# A combined neuroanatomy, ex vivo imaging and immunohistochemistry defined MRI mask for the human paraventricular nucleus of the thalamus

**DOI:** 10.1101/2025.04.29.651339

**Authors:** Madison R. Tetzlaff, Bianca T. Leonard, Michael A. Yassa, Tallie Z. Baram, Jerod M. Rasmussen

## Abstract

The paraventricular nucleus of the thalamus (PVT) is an evolutionarily conserved midline thalamic structure known to contribute to arousal, interoceptive states, and motivated behaviors. Yet, a consensus anatomical definition of the human PVT across tissue-based and MRI-based approaches remains elusive, thereby limiting reliable translation between its cellular characteristics and in vivo functional connectivity. To address this challenge, we describe a histologically-informed PVT segmentation compatible with standard 3T MR imaging pipelines. We performed postmortem anatomical MRI scans on an intact whole brain and an excised thalamic block, manually segmented the PVT at high resolution using ex vivo calretinin staining and neuroanatomical landmarks, registered the resulting image-label pair to a commonly used MRI template space (Montreal Neurological Institute’s MNI152), and performed a comparative reanalysis using this newly defined mask. This tissue-grounded PVT mask largely overlaps spatially with existing MRI-based PVT masks, with the exception of additional voxels posteriorly. Importantly, the functional connectivity patterns of this tissue-grounded mask are highly consistent with those previously reported. Collectively, this multimodal definition of the human PVT balances tissue-based ground truth with in vivo MRI features, providing a valuable resource for advancing translation between cellular level features identified by histology and in vivo functional connectivity at the meso/macro scale in the understudied human PVT.

**Key Points:**

- Addresses conflict between histological and MRI-based borders of the human PVT.
- Histologically-defined PVT boundaries indicate additional posterior tail to the human PVT as compared to existing human MRI atlases.
- Provided MRI mask demonstrates functional connectivity patterns consistent with prior human and rodent studies.

## Introduction

The paraventricular nucleus of the thalamus (PVT) is a midline thalamic nucleus that is highly conserved across species. It has been extensively studied in animal models ranging from lizards to rodents, underscoring its fundamental role in a wide array of behaviors (S. Bhatnagar et al. 2002; Seema Bhatnagar and Kirouac 2021; Fenoglio, Chen, and Baram 2006; Gao et al. 2023, 2020; Hain et al. 2022; Heredia et al. 2002; Kooiker, Birnie, and Baram 2021). Specifically, the PVT has been linked to arousal, hunger, salience detection, and the processing of internal states (Ye, Nunez, and Zhang 2022; Millan, Ong, and McNally 2017; Labouèbe et al. 2016; Engelke et al. 2021; Do-Monte, Quiñones-Laracuente, and Quirk 2015; Kooiker et al. 2023; Choi et al. 2019; Choi and McNally 2017; McNally 2021; Beas et al. 2024). The PVT is thought to play a critical role in regulating motivated behaviors, particularly by resolving conflicts between competing motivational drives. Despite its demonstrated importance, the human PVT remains relatively understudied, with a paucity of information regarding its cellular and chemical composition as well as its in vivo connectivity and functions.

Disagreement on the anatomical boundaries of the human PVT are a major barrier to its reliable study. Limited work directly examining the chemical and cellular composition of the human PVT include Uroz, et al, 2004 and Schulmann et al. 2024 (preprint), neither of which describe clear boundaries or cell type variations across the PVT. In contrast, the rodent PVT has been extensively examined by immunohistochemistry, spatial transcriptomics, and sequencing methods, and describes an important distinction between the anterior and posterior PVT (Gao et al. 2023; Shima et al. 2023; Beas et al. 2024; Kooiker, Birnie, and Baram 2021; Choi et al. 2019). The issue of anatomical boundaries in the human thalamus is not unique to the PVT; ambiguities in terminology and boundary definitions exist for all thalamic nuclei (Mai and Majtanik 2018). Several atlases have been created in attempt to unify these boundaries, though some group the PVT under the broader definition of ‘midline thalamus’, and others do not delineate the PVT at all, underscoring the extent to which its boundaries remain unsettled (Erzurumlu, Sengul, and Ulupinar 2024; Saranathan et al. 2021; Reeders et al. 2023; Pfefferbaum et al. 2023).

Recent in vivo human studies have relied on a putative three-dimensional thalamic atlas built from the Morel stereotactic atlas (Morel 2007; Krauth et al. 2010; Jakab et al. 2012; Kark et al. 2021), which includes PVT boundaries from eight post-mortem human brains. Using this atlas, work by several groups, including our own, suggests that human and rodent PVT networks share analogous target nodes, including the nucleus accumbens, amygdala, hypothalamus, ventral tegmental area, periaqueductal grey, and hippocampus (Li and Kirouac 2012; Otis et al. 2019; Kark et al. 2021; Kooiker, Birnie, and Baram 2021; McGinty and Otis 2020). Moreover, these connections appear behaviorally relevant, as they have been linked with prolonged maternal grief after loss of a child (Kark et al. 2022), anhedonia in adult males (Leonard et al. 2024), and craving in cocaine use disorder (Engeli et al. 2023). While these recent efforts with fMRI have advanced understanding of human PVT function, clear and mechanistic interpretation of these results is limited by the lack of clear, universally accepted neuroanatomical boundaries.

The Krauth MRI atlas (constructed using the Morel stereotactic atlas) represents the current state-of-the-science for fMRI of the human thalamus and serves as the foundation for most human PVT research, but the average-based delineation of nuclei may blur individual anatomical boundaries. For example, marked differences in PVT boundaries emerge when comparing the Morel-defined PVT borders with individual subject histology, such as that in the Allen Brain Atlas (Ding et al. 2016), the Paxinos atlas of the human brain (Mai, Assheuer, and Paxinos 2004), and the Nextbrain atlas (Casamitjana et al. 2024). The Morel and Krauth atlases appear to omit the posterior extent of the human PVT, but it is unclear if this represents an important or detectable variation in human anatomy and function. Thus, there is a pressing need for a histologically validated definition of the PVT to support ongoing in vivo human neuroimaging studies and interpretation of these studies in the context of cellular characteristics.

Here, we address anatomical inconsistencies in PVT delineation by constructing a “ground-truth” MRI-compatible three-dimensional mask based on histological markers from a single post-mortem human brain. We define PVT boundaries based on the positions of white matter tracts and the continuous, circumscribed expression of calretinin, an established marker of the PVT across multiple species including human, primate, cat, rat, mouse, and lizard (Fortin, Asselin, and Parent 1996; Gao et al. 2023; Hua et al. 2018; Uroz, Prensa, and Giménez-Amaya 2004; Winsky et al. 1992). This combined approach identifies a set of boundaries for the PVT common across postmortem histology, neuroanatomy and MRI-visible features. In addition, we reconcile discrepancies with the commonly used Morel atlas through a descriptive analysis of these boundaries and a reanalysis of MRI-based in vivo functional connectivity data.

## Methods

The overall design and workflow of the experiments is shown in Figure 1. Briefly, the postmortem brain was scanned at 3T, and then thalamus and overlying ventricular system were dissected out to a block and re-scanned at both 3T and 9.4T. After completion of MRI scans, the thalamic block was further processed into sections, stained for calretinin, and the PVT was traced back onto the highest resolution MRI scan. Finally, the resulting PVT mask was transformed into MNI standard space and the functional connectivity was assessed as compared to that of our previously published MRI mask (Kark et al, 2021). A more detailed description of these procedures is described below.

**Figure 1.**
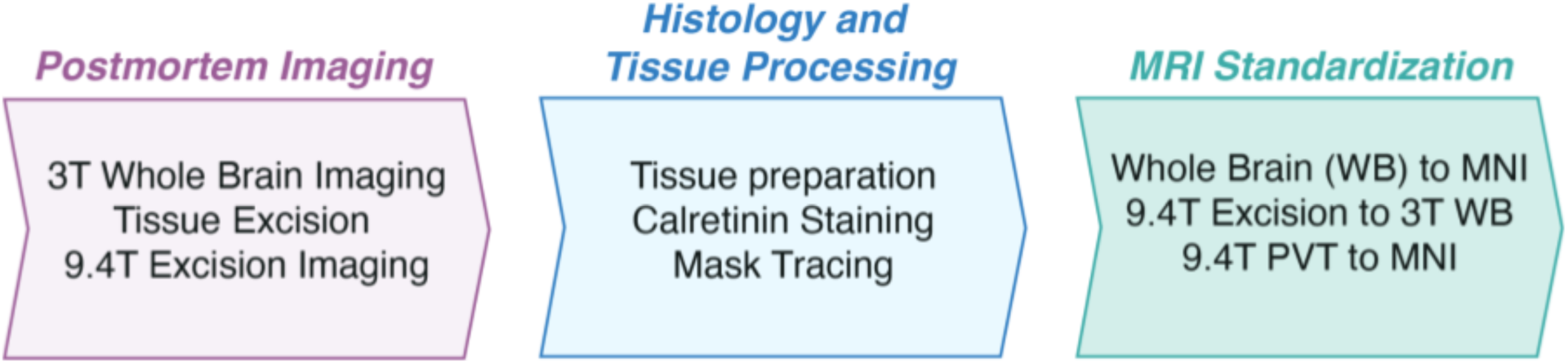
Overview of experimental framework. Postmortem MRI scans were acquired at two different resolutions, before and after the excision of the bilateral thalamic region (“thalamic block”). Histological processing followed, including calretinin staining for enhanced visualization of the PVT region. Guided by the histology results, a three-dimensional PVT mask was segmented. Through linear and nonlinear transformations, this mask was rendered in standard MNI152 template space for functional connectivity analysis using fMRI.

### Post-mortem MRI

A 67-year-old male with no known neuropsychiatric or neurodegenerative disease contributed his brain to the UCI Willed Body program and formed the basis of the procedures and analyses performed here.

Prior to histological processing, we acquired a series of high-resolution MR images of the whole postmortem brain to facilitate the creation of a histologically grounded PVT mask in standard template space. Each acquisition was designed to optimize registration and anatomical fidelity across modalities. First, we obtained a standard and high-resolution whole-brain T1-weighted image at 3T to maximize intensity and contrast information for registration to the MNI template. The brain was then dissected to remove cortical structures, leaving the ventricular system largely intact (Supplemental Fig. 1), and re-scanned at 3T using identical sequences to capture image details while accounting for deformations introduced by the dissection. Finally, the thalamic block was scanned at 9.4T with ultra-high resolution ensuring precise anatomical correspondence with the histological images. Collectively, these imaging steps serve as intermediate reference points, maximizing information overlap and ensuring accurate registration between histology and the commonly used MNI152 MRI template. Specific acquisition details are provided below.

### Whole Brain and Excised (“thalamic block”) 3T MRI Acquisition

Postmortem MRI imaging was conducted on a 3T Siemens MAGNETOM Prisma scanner. The imaging protocol included a high resolution structural MPRAGE sequence (0.8 mm isotropic resolution; 320 sagittal slices, field of view = 240 mm × 256 mm, flip angle = 4, TR/TE = 5000/3.2 ms, matrix size = 300 × 176, inversion pulse TI = 700 ms). Following whole brain imaging, the thalamic block, which contained the PVT and adjacent structures was immersed in fluorinert and re-scanned at 3T using identical imaging parameters to maintain consistency while accounting for potential post-processing deformations.

### Excised 9.4T MRI Acquisition

The block was then rescanned using a 9.4 Tesla (9.4T) Bruker scanner to obtain ultra-high resolution structural imaging. A spin echo sequence was used (0.6 mm isotropic resolution; 224 sagittal slices, field of view = 200 × 200 mm, TR/TE = 1100/3.2 ms, matrix size = 320 × 320).

### Histology and Tissue Processing

Following the 9.4T scan, the excised block was submerged in 15% sucrose in 0.1M phosphate buffer followed by 30% sucrose in phosphate buffer, allowing the tissue to sink in each solution to ensure proper water removal and cryoprotection. The tissue was then trimmed and sectioned into 1.5 cm coronal slices for ease of handling, embedded in OCT *(Tissue-Tek*, *#4583*) in flat trays (Supplemental Fig. 1), and snap-frozen in a bath of isopentane chilled by dry ice. The frozen tissue was sectioned at 40 microns using a Leica CM1900 cryostat and stored in 0.1M phosphate buffered saline (PBS) with 0.01% sodium azide at 4°C.

Due to prolonged fixation (>3 months) in 10% formalin, formalin stripping was required to unmask the calretinin epitope. This was accomplished via incubations in 1M EDTA (pH 8.0) and 1x Triton, heated to near boiling and maintained at this temperature via microwave bursts (∼10 seconds each) for 20 minutes. Following formalin stripping, the specimen was rinsed in 1x PBS with Triton (PBST), treated with 30μL/10mL of 30% H_2_O_2_ for 30 minutes, then blocked for 1 hour in 10% normal goat serum (NGS). Primary antibody incubation for 36-48 hours at 4°C with polyclonal rabbit anti-calretinin (1:250, *Invitrogen, #180211*) in 2% NGS in 1x PBST was followed by secondary incubation for 2 hours at room temperature in biotinylated goat anti-rabbit (*Invitrogen, #3182G)* in 1x PBST. After antibody incubations, tissue was incubated with ABC-Kit (*Vector Labs, PK-4000*) for 2-3 hours per manufacturer instructions. Tissue was allowed to react with activated DAB (*Vector Labs, SK-4100*) for 12-15 minutes before ceasing the reaction. When used, hematoxylin (*Abcam, AB220365*) was applied for 30 seconds before rinsing with 1x PBST. Microscopy images were taken with a Nikon Eclipse E4000 microscope with the use of the Nikon FS Elements acquisition software. Gross anatomical images were taken with an iPhone 13.

### Alignment of the tissue specimen with Post-mortem MRI

Gross anatomical markers including the mammillary bodies, red nucleus, mammillothalamic tract, habenular commissure, other white matter tracts, and the ventricular contours were used as landmarks to align the tissue specimen with MRI sections. Calretinin-positive regions of midline thalamus were manually segmented over coronal sections of the 9.4T excised thalamic MRI image (0.2 mm isotropic) to generate a 3D seed region using 3D Slicer (v5.6.2).

The first step in generating a transformation matrix to place the PVT segmentation in standard space involved aligning the histological tracing on the 9.4T image (0.2 mm resolution) to the 0.6 mm resolution image from the same scanner using linear transformation (Fig. 4, Step 1). This was performed using FMRIB’s Linear Image Registration Tool, FLIRT, which is part of the FMRIB Software Library (FSL; Jenkinson et al. 2012).

### Alignment of 9.4T Thalamic Block to 3T Postmortem Whole Brain

To generate an artificial excision from the successfully registered 3T whole-brain scan, the 9.4T image of the thalamic block was registered to the 3T image of the same block (Fig. 4, Step 2). A binary mask of the 3T thalamic block was generated within the whole brain to support the registration of the 3T thalamic block to the 3T whole brain image (Fig. 4, Step 3,4). Finally, the 3T thalamic block and whole-brain images were registered using the nonlinear transformation matrix generated in the next step, aligning the 3T whole-brain scan to MNI152 standard space (Fig. 4, Step 5).

### Alignment of 3T Postmortem Whole Brain to MNI152 Standard

To facilitate the creation of an MRI atlas in standardized space, we performed brain extraction (Brain Extraction Tool, BET) and linear registration of the 3T postmortem whole-brain scan. A nonlinear transformation (12 DOF) was then applied using Advanced Normalization Tools (ANTs; Avants, Tustison, and Song 2009) to align the scan with the MNI152 brain template at 0.5 mm resolution (Fig. 4, Step 6).

### Alignment of the histology Informed PVT Segmentation to MNI152 Standard

The transformation matrices from the previous section were concatenated and applied to the histology-informed PVT mask using FLIRT (Fig. 4, Step 7). The resulting voxels were thresholded at 0.5, after which the mask was binarized. This final step completed the transformation of the PVT segmentation from the native 9.4T space into an MNI152 standard space PVT mask with a 0.5 mm resolution. To assess consistency, a Dice-Sørenson coefficient (DSC) was calculated between the previous and updated PVT masks.

### Statistical analysis of variation in masks and functional connectivity

A critical comparison between the histologically-defined PVT mask and the commonly used Morel atlas PVT mask revealed notable differences in anatomical definition. To assess the functional implications of these differences, we compared functional connectivity measures derived from the new histologically defined PVT mask with those derived from the Morel atlas PVT mask. This re-analysis was conducted using previously published PVT connectivity maps (Kark et al. 2021) ensuring that all methodological aspects remained identical, except for the mask used.

The imaging sample for this functional connectivity analysis was drawn from the Human Connectome Project (HCP) young adult cohort. It included 121 young healthy adults (ages 22-35, 73 female, 48 male) who underwent a 7T resting state functional magnetic resonance imaging (rsfMRI) session. This sample comprised all remaining participants after exclusions for poor image quality, excessive motion in the scanner, or anatomical, behavioral, or neurological criteria flagged by the HCP protocol.

Full details of the image processing and functional connectivity analyses are available in our group’s previous study (Kark et al. 2021). Briefly, we used the ‘‘minimally preprocessed’’ datasets provided by the HCP 1200 Release (HCP filename: rfMRI∗hp2000_clean.nii.gz, n = 135) (Glasser et al. 2013). These datasets included structural and functional images that had undergone preprocessing, including independent components analysis for artifact removal of noise components from fMRI data (ICA-FIX) (Salimi-Khorshidi et al. 2014; Griffanti et al. 2014). Consistent with Kark et al., both structural and functional datasets were further processed using the CONN Toolbox for functional connectivity analyses (Smith and Nichols 2009; Whitfield-Gabrieli and Nieto-Castanon 2012). Within the CONN toolbox, data were segmented, smoothed, denoised, and processed for further artifact detection via Artifact Detection Tools (ART), where conservative motion censoring thresholds were applied, and 14 of the 135 participants were dropped for having fewer valid scans than the rest of the group (as defined by 1st Q - 1.5 IQR).

Functional connectivity seed-to-voxel maps were then generated using the new PVT segmentation. Connectivity was calculated by calculating semipartial correlations between the average BOLD signal of the PVT mask voxels and all other voxels in the brain. The PVT mask was subtracted from a whole-thalamus mask (Jakab et al. 2012), and the signal from the remaining thalamic voxels were added as a regressor to the calculation of functional connectivity for the PVT. Given the small volume of the PVT mask, this was done to reduce signal from overlapping voxels to the PVT functional connectivity calculation. To determine significant connectivity patterns, we applied a threshold-free cluster enhancement (TFCE) method with family-wise error correction (pTFCE-FWE<0.05) with 1,000 permutations. The degree of overlap between spatial maps was quantified using DSC. Coefficient of determination (*R*^2^) values were calculated from T-statistic maps of the functional connectivity measures to further quantify the similarity between each seed-to-voxel functional connectivity map.

## Results

### Tissue-based definition of the human PVT

Histologically, the PVT was defined as a calretinin-positive, continuous region bordering the third ventricle and ventral to the stria medullaris thalami. In the coronal plane, its widest point measured 800 micrometers at the level of the mammillothalamic tract, while its narrowest point measured approximately 200 micrometers, extending posteriorly bordering the mediodorsal nucleus and the third ventricle (Fig. 2). As the PVT approached the anterior region of the habenular commissure, it widened slightly to about 600 micrometers. Posterior to the interthalamic adhesion and ventral to the habenula, calretinin staining revealed an abundance of fibers extending dorsolateral to the third ventricle, in what is often referred to as the ”periventricular area” (Fig. 2E). Given the distinct change in calretinin signal type in this region, we determined that these fibers were outside of the PVT.

**Figure 2.**
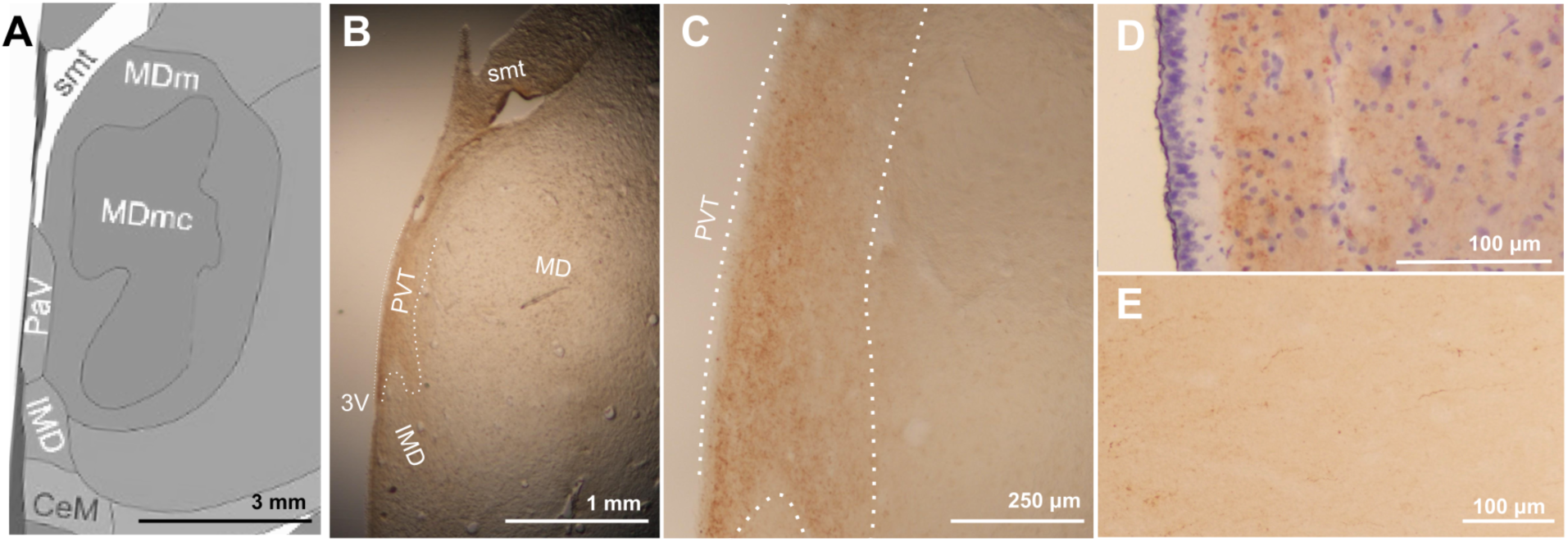
Calretinin delineates paraventricular nucleus of the thalamus from neighboring nuclei. (A) Representative region from Allen Human Brain Reference Altas (*PaV = PVT)* and (B) corresponding brightfield image of human midline thalamus stained for calretinin with the PVT outlined along calretinin borders used to create MRI mask. Cellular puncta are visible at higher magnification of the defined area (C) and hematoxylin and calretinin stain show specificity of calretinin labeling away from ependymal layer (D). Past the posterior extent of the PVT, calretinin stain defines diffuse fibers (E). Abbreviations: 3V: Third ventricle, CeM: Centromedial nucleus, IMD: intermediodorsal nucleus, MD: Mediodorsal nucleus, MDm: Mediodorsal nucleus, medial part, MDmc: Mediodorsal nucleus, magnocellular part, PVT: paraventricular nucleus of the thalamus, smt: stria medullaris thalamica

### Alignment of histologically-defined PVT with MRI images

The calretinin- and neuroanatomy-defined PVT region was traced on the highest resolution MRI available for the post-mortem excised thalamus block at 9.4T (Fig. 3a). The resulting three-dimensional shape of the bilateral PVT was largely symmetrical, with a contour that closely matched the anatomical PVT and aligned with the tissue-based observations above. Specifically, both left and right PVT were relatively wide anteriorly, narrowed along the extent of the mediodorsal thalamus, and exhibited a slight ventral dip (Fig. 3b). The initial histologically grounded volume was approximately 63 mm^3^, encompassing 3,148 voxels at 0.2 mm resolution. Following smoothing and registration to standard space at a lower, 0.5 mm voxel resolution, the new segmentation contained 1,583 voxels (∼198 mm^3^). In standard space, a critical comparison between this histologically grounded mask and the previously published Morel atlas-based mask revealed a substantial degree of overlap in the anterior region, but a limited global overlap (DSC = 0.37). The histologically grounded mask also featured an additional posterior extension not present in the Morel atlas-derived mask, which appears to contribute to the majority of non-overlapping voxels (Fig. 3c).

**Figure 3.**
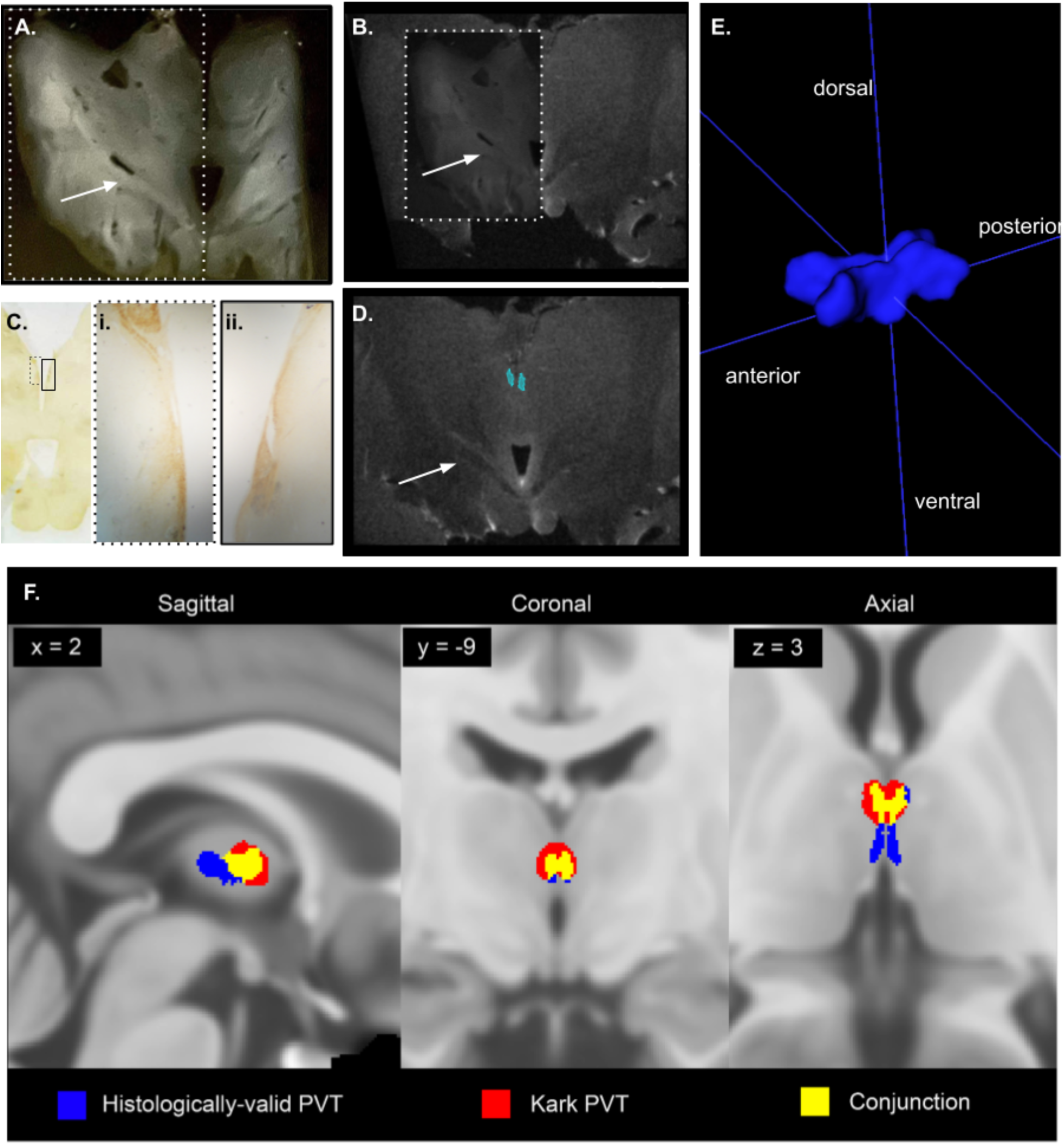
Alignment of histology to MRI produces posterior tail on PVT mask. Example alignment of tissue slice to MRI section, showing wet, unstained slice with distinguished white matter tracts (A) aligned to MRI slice in (B). White dotted border in (A) outlines part of image which is made 50% opaque and overlaid on (B). Calretinin-stained slide (C) neighboring (A) with Ci. left and Cii. right boxes showing the calretinin positive region traced onto the MRI slide as in (D). Arrow distinguishes common white matter tract in same location across slides. (E) Three-dimensional rendering of final PVT mask in MNI152 standard MRI space (volume = 1,583 voxels (∼198 mm3). (F) Comparisons between the PVT mask employed in the previous functional connectivity study (red; Kark, 2021) and the newly generated PVT mask (blue; Dice-Sørenson coefficient (DSC) = 0.37).

### Functional connectivity of the histologically validated human PVT mask

To assess the practical implications of the current and previous PVT masks in the context of in vivo human fMRI, we compared functional connectivity (FC) maps generated using the histologically grounded mask with the FC maps previously published using the Morel atlas. Overall, the two masks produced highly congruent whole-brain connectivity patterns (Fig. 4) with strong agreement in both positive FC maps (DSC = 0.854, *R*^2^ = 0.851) and negative FC maps (DSC = 0.848, *R*^2^= 0.857). Importantly, as observed in prior studies using Morel atlas-based masks, the functional connectivity of the human PVT remained consistent with findings from rodent studies. Regardless of the mask used, the human PVT exhibited strong connectivity with key nodes of both the reward and default networks, reinforcing cross-species convergence in its functional organization.

**Figure 4.**
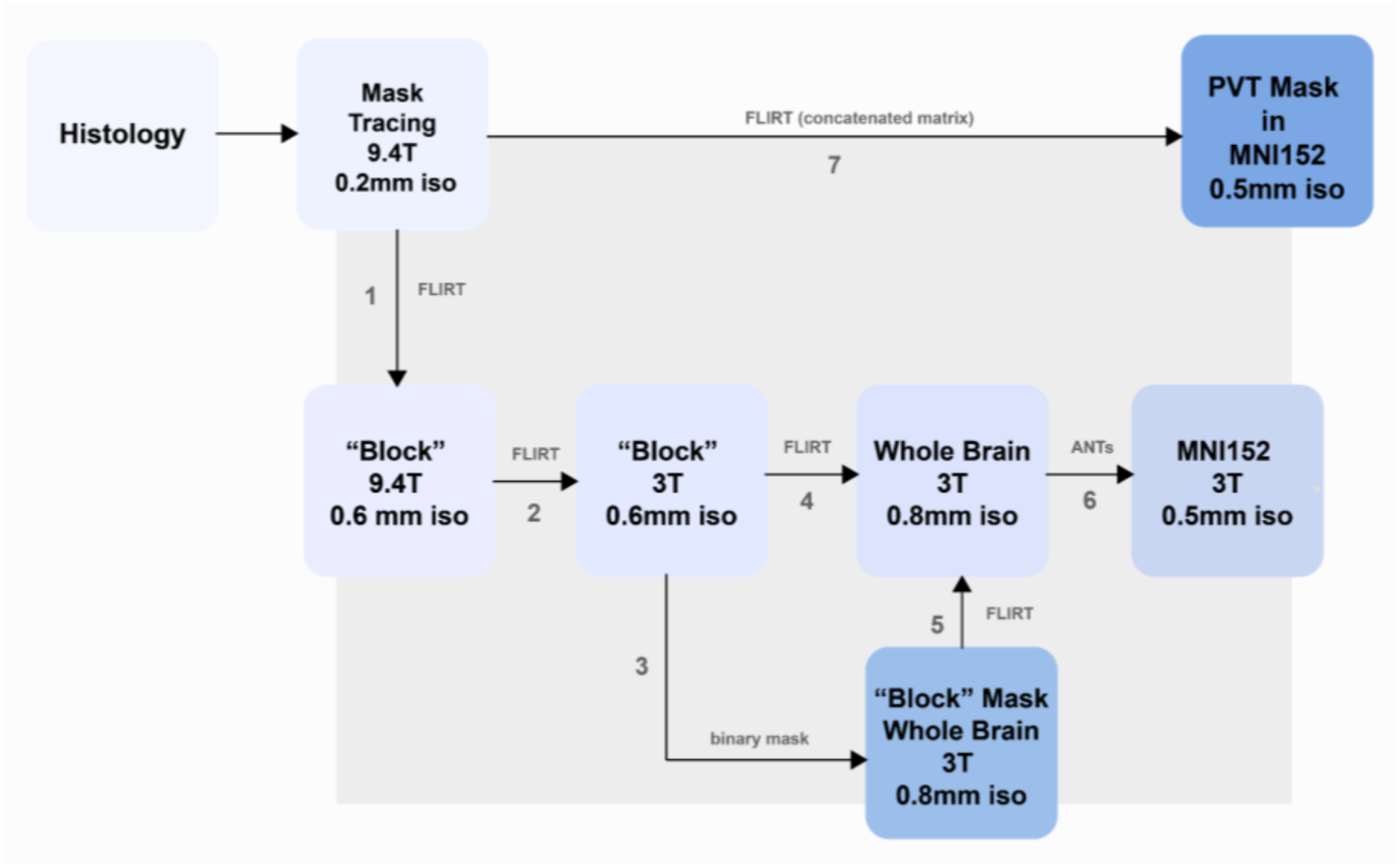
Registration pipeline to transform PVT mask from histological-tracing space into MRI standard space (MNI152). Several structural images were captured of both the whole postmortem brain and an excision of the thalamus and surrounding tissues (“block”) at differing resolutions and by leveraging different scanners to aid in the transformations at intermediate steps for improved registration quality of the final PVT mask.

**Figure 5.**
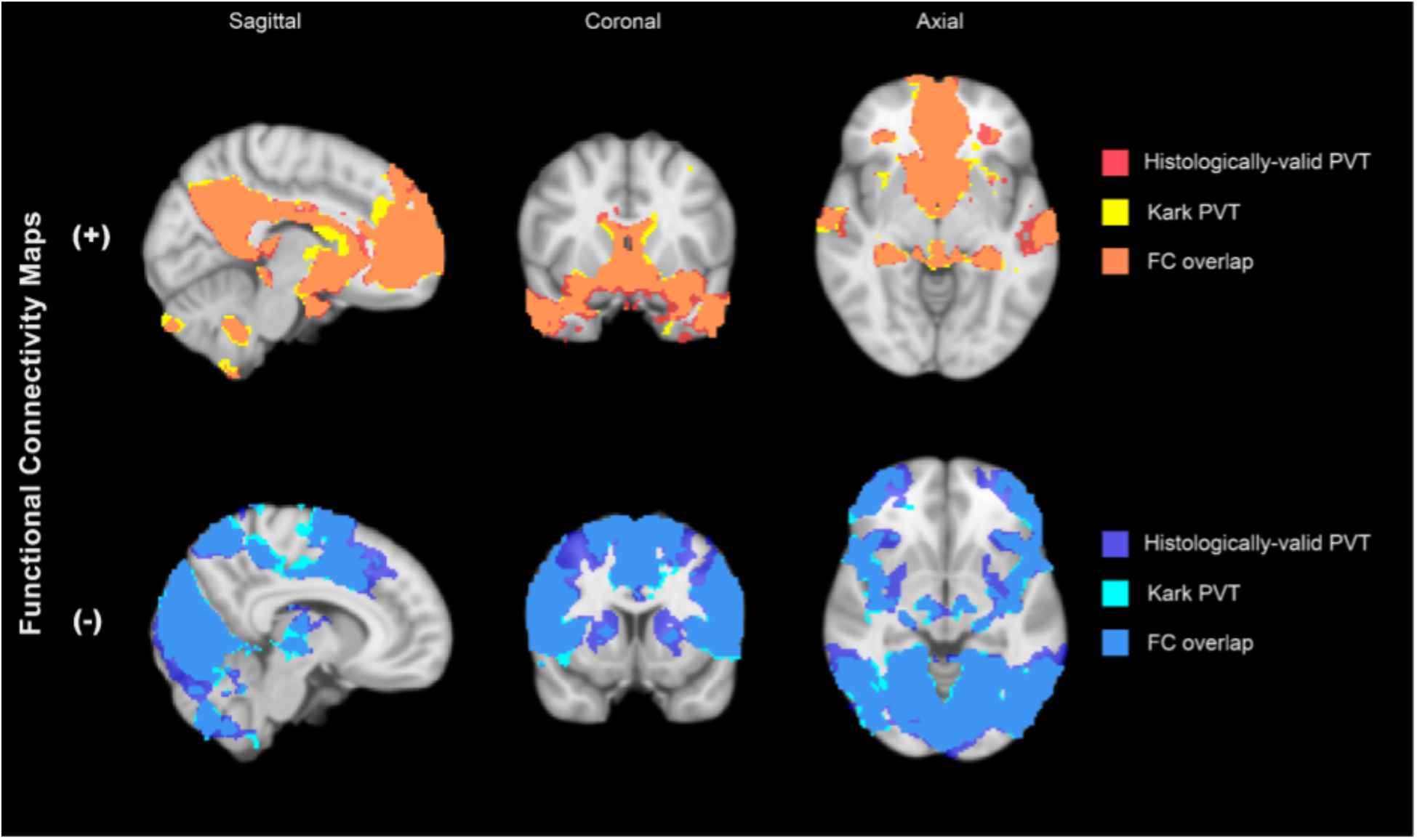
Comparative functional connectivity between histology-based mask and Kark et al. 2021. Functional connectivity (FC) of PVT is highly similar between the Kark et al. 2021 segmentation, and the histologically valid segmentation presented in this study. Following threshold-free cluster enhancement (pTFCE-FWE <0.05) selection of significant voxels, the maps from each analysis were divided by the sign (+/-) of the functional connectivity measures, and compared by calculating the Dice-Sørenson coefficients (DSC) and coefficients of determination (R2) to quantify similarity between maps (positive FC maps: DSC = 0.854, R2 = 0.851; negative FC maps: DSC = 0.848, R2= 0.857).

## Discussion

The key contributions of this study address challenges related to the translation across tissue-based and MRI-based definitions of the PVT. First, by integrating neuroanatomical landmarks with cell-type-specific immunohistochemistry, informed by modern atlases, we delineate the three-dimensional boundaries of the human PVT with greater specificity than previously available. Second, we reanalyzed MRI-based in vivo functional connectivity using this new mask and found it to be highly consistent with our previously published findings, thereby validating both the new tissue-grounded masks and existing approaches as useful tools for studying this critical, yet understudied brain region. Finally, we provide the human neuroimaging community with the first segmentation of the human PVT derived directly from postmortem tissue and transformed into a commonly used MNI152 template space, allowing for increased interpretation between cellular characteristics and large-scale in vivo brain networks.

### Integrating Tissue-based and MRI-based Approaches Aids the Study of the Human PVT

Calretinin staining was used to distinguish the PVT from neighboring nuclei in postmortem tissue sections. Calretinin is an established marker of PVT across species; however, while abundant in the PVT, it lacks specificity relative to the thalamic paratenial and reuniens nuclei, as well as the centromedial nucleus and the habenula. Nonetheless, these neighboring nuclei can be readily distinguished in coronal human tissue sections based on their proximity to the ventricular edge and position along the dorsoventral axis. We therefore delineated the PVT from the calretinin-negative mediodorsal thalamic nucleus using calretinin, and from other calretinin-positive nuclei using anatomical landmarks to produce a ground-truth postmortem tissue definition of the human PVT.

In contrast to postmortem microscopy, the boundaries of small thalamic structures such as the PVT, paratenial nucleus, and reuniens nucleus are not easily distinguishable on standard-resolution structural MRI. This limitation to precise localization necessitates the multimodal approach used here, while also requiring anatomical landmarks for cross-modal integration. To ground our translation, we relied on myelinated structures that are clearly identifiable on both sectioned tissue and MRI, including the mammillothalamic tract and the anterior and habenular commissures. Based on our observations, these structures serve as reliable landmarks for anchoring masks of the PVT and other small midline thalamic nuclei within the neuroanatomy visible in MRI space (MNI152). Thus, future efforts to individualize MRI-based segmentations of small, variable midline thalamic nuclei may benefit from using these landmarks as grounding points for transformation to standard space.

We observed substantial volumetric differences in the PVT when transforming from native space at ultra high-resolution (0.2 mm isotropic at 7T) to template space at high-resolution (0.5 mm isotropic at 3T). While this can largely be attributed to the spatial smoothing and quantization inherent in registration and down-sampling processes, it also highlights challenges associated with translating small laminar structures to relatively low resolution spaces like those used in fMRI analysis. MRI-based technologies continuously drift towards higher resolution acquisition, but at current resolutions this inflation is likely to remain, particularly when used for fMRI. A single, modern fMRI voxel is roughly 2 mm in width, wider than the widest coronal point of the PVT, suggesting that the partial sampling of surrounding nuclei is likely. This concern can be addressed in functional connectivity studies by regressing out the BOLD signal of voxels near PVT boundaries, as done previously by our group (Kark et al. 2021). We suggest that those investigating either the histological or functional human PVT should remain conscious of the atlases used to define the PVT in relation to the technology they implement.

As future work to study the PVT with either post-mortem ex vivo or fMRI in vivo methodologies continues, our work provides a shared ground which could prove essential to comparative or correlative efforts across modalities. Specifically, appropriate comparison across these technologies allows consideration of the in-vivo-derived functional significance of any ex-vivo-derived cellular variations across the anterior-posterior axis of the human PVT. We anticipate that recent advances in post-mortem oriented technologies such as spatial transcriptomics are likely to dramatically improve our knowledge of small, variable human brain structures by allowing groups to generate high-dimensional data with unprecedented anatomical precision. As this post-mortem body of data grows, we believe that studies such as ours will serve as an essential bridge between functional, in vivo phenomenon and the underlying cellular encoding processes.

### Tissue-based Definitions of the PVT Extend Posterior Compared to Existing Available Atlases

No clear distinction between the anterior and posterior human PVT was observed in this study. Notably, this is in contrast to reports across species of an anterior and posterior division to the PVT. Specifically, rodent studies indicate functional and cellular differences across the anterior-posterior axis, with the anterior PVT associated with arousal and reward behaviors and the posterior PVT involved in stress and fear responses. Selective activity along this axis has been reported during motivated behavior (Beas et al. 2024) and as a result of early-life stress (Kooiker 2023). In humans, an anatomical division characterized by a thin laminar strip connecting the anterior and posterior PVT was recognized long before functional or cellular distinctions were characterized in any species. In terms of calretinin expression, we found no clear distinction between the anterior and posterior human PVT. However, the mid-PVT narrowing we report is consistent with previous work dividing the human PVT into anterior and posterior segments, and could represent a natural point of division. Toward the posterior end, the PVT gradually widens, with decreasing cell density that transitions into fiber tracts in the periventricular area (Dewulf 1971; Mai, Assheuer, and Paxinos 2004; Van Buren and Borke 1973). Thus, we propose that these fiber tracts are not within the PVT, but instead constitute the posterior boundary. When comparing our histologically guided definition of the PVT to existing atlases in MRI space, we observed that this posterior extension is largely not captured by such atlases, and represents an important area of future investigation.

Diverse perspectives exist regarding the extent and boundaries of the human posterior PVT. An additional region, confusingly referred to as the “periventricular area of the thalamus” (PeVa) is sometimes included as the posterior end of the PVT (Ding et al. 2016); in this region, we found calretinin-positive fibers, but no cell bodies, supporting the conclusion that this “PeVa” region is not part of the paraventricular *nucleus* of the thalamus. Notably, the terminology “paraventricular thalamus” is sometimes also used to describe the midline nuclei bordering the ventricle in the human; this likely contributes to confusion regarding the PVT and its borders, and we hope this work serves as clarification of the entity that is the human paraventricular *nucleus* of the thalamus, referred to here as the PVT.

### Functional Connectivity Maps With and Without the Posterior End of the PVT are Highly Similar

We present a new anatomically and histologically informed segmentation of the human PVT for use in in vivo human functional connectivity studies. Because this mask extends posteriorly relative to previous definitions, we compared functional connectivity maps derived from the new segmentation to those from our prior mask based on existing atlases. Notably, the inclusion of the posterior segment appeared to have minimal impact on the detection of functionally connected nodes using high spatial resolution fMRI data from the 7T HCP Project. The high degree of overlap in functional connectivity between the two masks, along with their agreement with rodent connectivity patterns, supports the notion that human and rodent PVT networks target analogous regions. Given that the new mask spans both anterior and posterior portions of the PVT, and the previous version included primarily anterior PVT, our functional connectivity findings are consistent with the absence of histological differences on the posterior-anterior axis as revealed by calretinin staining. However, this result may be alternatively explained by the relatively large point-spread function in fMRI, combined with the spatial extent and vascularization of the PVT, which may obscure fine-grained functional distinctions (Fracasso, Dumoulin, and Petridou 2021). Future studies leveraging higher-resolution imaging and targeted functional analyses may reveal anterior-posterior specialization within the human PVT, with relevance for both normative and clinical populations.

While the current study was designed to address challenges in interpreting between studies performed at the cellular and meso/macroscopic scale, limitations remain—particularly regarding interindividual variability in borders not easily observed at current standard MRI resolutions. To help address this, we compared our tissue-based segmentation, derived from a single individual, with existing prints of both individual and averaged post-mortem thalamic parcellations. Based on qualitative comparisons, our donor-derived PVT aligns with that of five postmortem cases used in the construction of the NextBrain atlas (Casamitjana et al. 2024), as well as with other established atlases (Mai, Assheuer, and Paxinos 2004; Toncray and Krieg 1946). Specifically, we found that our PVT parcellation and others always begin the PVT slightly anterior to the mammillothalamic tract, PVT thins into a laminar strip as it proceeds posteriorly, and that the posterior edge of the PVT is at the level of the habenulae. Notably, some atlases show a slightly wider PVT anteriorly, and a slightly narrower PVT posteriorly as compared to our individual; these variations are on the scale of less than half a millimeter, and may represent natural individual variation. The general agreement of our individual PVT with over 30 cumulative other reported cases of individual thalamic parcellations supports the accuracy of our delineation and suggests that our donor does not represent a gross anatomical outlier.

Nonetheless, substantial variation does exist in the human midline thalamus, including differences as obvious as the presence or absence of the interthalamic adhesion (massa intermedia) and gross branching of vascularization affecting the PVT region (Castaigne et al. 1981; Damle et al. 2017; Percheron 1973; Şahin et al. 2023; Vidal et al. 2024; Kochanski et al. 2018). Across the 25 cases averaged by Van Buren and Borke (1973), variation in the reported boundary for mammillothalamic tract exceeded 5 mm, which would span multiple voxels at standard MRI resolutions. Given this variability, it is plausible that the averaged, anteriorly-biased PVT present in the Morel and Krauth atlases represents a consistent “core” of the human PVT. Our findings extend this notion by demonstrating that functional connectivity derived from both our tissue-based and the atlas-based masks exhibit substantial overlap, supporting their shared utility in characterizing PVT.

Recent progress leveraging high-resolution fMRI data has revealed brain-wide functional connectivity patterns of the human PVT (Kark et al. 2021). The current study builds on this work by incorporating a histologically grounded definition of the human PVT. This newly developed mask is now publicly available in MNI152 space, ensuring broad accessibility for neuroimaging research. While the observations made here suggest that masks currently in popular use (*i.e.,* Krauth) remain suitable for use in standard MR image processing pipelines and may reflect a “core” region of the PVT likely to be present across individuals, this newly available mask may provide a more comprehensive PVT including an additional posterior region. The use of this new mask may provide further validation and new insights into the function of the human PVT in health and disease.

## Conclusion

In summary, the current work provides the neuroimaging community with a valuable new resource: the first critical comparison between an anatomically and cell-type informed segmentation of the human PVT, and an average-based mask of the PVT. It reconciles differences in extant atlases and affirms the validity of prior masks and their utility in analyses of PVT functional connectivity. We anticipate that this resource will become a useful tool to support future research into the structure and function of the human PVT, an important, yet understudied brain region.

## Acknowledgements

The authors wish to thank individuals who donate their bodies and tissues for the advancement of education and research.

We are grateful for the technical assistance and advice from both Drs. Andre Obenaus and Nick Tustison regarding the acquisition of 9.4T MR images and MR image registration, respectively.

Funding was provided by P50 MH-096889 (PI: TZB), R01 MH-132680 (PI: TZB), R01 MH-128306 (PI: MAY), T32 MH-119049 (PI: MAY, BL), R00 HD-100593 (PI: JMR), T32 GM008620/GM/NIGMS (BL, MT), and the Bren Foundation.

## References

Avants, B., N. Tustison, and G. Song. 2009. “Advanced Normalization Tools: V1.0.” The Insight Journal, July. 10.54294/uvnhin.

Beas, Sofia, Isbah Khan, Claire Gao, Gabriel Loewinger, Emma Macdonald, Alison Bashford, Shakira Rodriguez-Gonzalez, Francisco Pereira, and Mario A. Penzo. 2024. “Dissociable Encoding of Motivated Behavior by Parallel Thalamo-Striatal Projections.” Current Biology: CB 34 (7): 1549–60.e3.

Bhatnagar, Seema, and Gilbert J. Kirouac. 2021. “Editorial: Advances in Understanding of the Functions of the Paraventricular Thalamic Nucleus.” Frontiers in Integrative Neuroscience 15 (August):744147.

Bhatnagar, S., R. Huber, N. Nowak, and P. Trotter. 2002. “Lesions of the Posterior Paraventricular Thalamus Block Habituation of Hypothalamic-Pituitary-Adrenal Responses to Repeated Restraint: Paraventricular Thalamus Lesions Block Habituation to Repeated Stress.” Journal of Neuroendocrinology 14 (5): 403–10.

Casamitjana, Adrià, Matteo Mancini, Eleanor Robinson, Loïc Peter, Roberto Annunziata, Juri Althonayan, Shauna Crampsie, et al. 2024. “A next-Generation, Histological Atlas of the Human Brain and Its Application to Automated Brain MRI Segmentation.” Neuroscience. bioRxiv. https://www.biorxiv.org/content/10.1101/2024.02.05.579016v1.full.pdf.

Castaigne, P., F. Lhermitte, A. Buge, R. Escourolle, J. J. Hauw, and O. Lyon-Caen. 1981. “Paramedian Thalamic and Midbrain Infarct: Clinical and Neuropathological Study.” Annals of Neurology 10 (2): 127–48.

Choi, Eun A., Philip Jean-Richard-Dit-Bressel, Colin W. G. Clifford, and Gavan P. McNally. 2019. “Paraventricular Thalamus Controls Behavior during Motivational Conflict.” The Journal of Neuroscience: The Official Journal of the Society for Neuroscience 39 (25): 4945–58.

Choi, Eun A., and Gavan P. McNally. 2017. “Paraventricular Thalamus Balances Danger and Reward.” The Journal of Neuroscience: The Official Journal of the Society for Neuroscience 37 (11): 3018–29.

Damle, Nishad R., Toshikazu Ikuta, Majnu John, Bart D. Peters, Pamela DeRosse, Anil K. Malhotra, and Philip R. Szeszko. 2017. “Relationship among Interthalamic Adhesion Size, Thalamic Anatomy and Neuropsychological Functions in Healthy Volunteers.” Brain Structure & Function 222 (5): 2183–92.

Dewulf, André. 1971. Anatomy of the Normal Human Thalamus: Topometry and Standardized Nomenclature / [by] A. Dewulf. Amsterdam: Elsevier Pub. Co.

Ding, Song-Lin, Joshua J. Royall, Susan M. Sunkin, Lydia Ng, Benjamin A. C. Facer, Phil Lesnar, Angie Guillozet-Bongaarts, et al. 2016. “Comprehensive Cellular-Resolution Atlas of the Adult Human Brain.” The Journal of Comparative Neurology 524 (16): 3127–3481.

Do-Monte, Fabricio H., Kelvin Quiñones-Laracuente, and Gregory J. Quirk. 2015. “A Temporal Shift in the Circuits Mediating Retrieval of Fear Memory.” Nature 519 (7544): 460–63.

Engeli, Etna J. E., Andrea G. Russo, Sara Ponticorvo, Niklaus Zoelch, Andreas Hock, Lea M. Hulka, Matthias Kirschner, et al. 2023. “Accumbal-Thalamic Connectivity and Associated Glutamate Alterations in Human Cocaine Craving: A State-Dependent Rs-fMRI and 1H-MRS Study.” NeuroImage. Clinical 39 (103490): 103490.

Engelke, D. S., X. O. Zhang, J. J. O’Malley, J. A. Fernandez-Leon, S. Li, G. J. Kirouac, M. Beierlein, and F. H. Do-Monte. 2021. “A Hypothalamic-Thalamostriatal Circuit That Controls Approach-Avoidance Conflict in Rats.” Nature Communications 12 (1): 1–19.

Erzurumlu, Reha, Gulgun Sengul, and Emel Ulupinar. 2024. Human Neuroanatomy. San Diego, CA: Academic Press.

Fenoglio, Kristina A., Yuncai Chen, and Tallie Z. Baram. 2006. “Neuroplasticity of the Hypothalamic-Pituitary-Adrenal Axis Early in Life Requires Recurrent Recruitment of Stress-Regulating Brain Regions.” The Journal of Neuroscience: The Official Journal of the Society for Neuroscience 26 (9): 2434–42.

Fortin, M., M. C. Asselin, and A. Parent. 1996. “Calretinin Immunoreactivity in the Thalamus of the Squirrel Monkey.” Journal of Chemical Neuroanatomy 10 (2): 101–17.

Fracasso, Alessio, Serge O. Dumoulin, and Natalia Petridou. 2021. “Point-Spread Function of the BOLD Response across Columns and Cortical Depth in Human Extra-Striate Cortex.” Progress in Neurobiology 202 (102034): 102034.

Gao, Claire, Chiraag A. Gohel, Yan Leng, Jun Ma, David Goldman, Ariel J. Levine, and Mario A. Penzo. 2023. “Molecular and Spatial Profiling of the Paraventricular Nucleus of the Thalamus.” eLife 12 (March). 10.7554/eLife.81818.

Gao, Claire, Yan Leng, Jun Ma, Victoria Rooke, Shakira Rodriguez-Gonzalez, Charu Ramakrishnan, Karl Deisseroth, and Mario A. Penzo. 2020. “Two Genetically, Anatomically and Functionally Distinct Cell Types Segregate across Anteroposterior Axis of Paraventricular Thalamus.” Nature Neuroscience 23 (2): 217–28.

Glasser, Matthew F., Stamatios N. Sotiropoulos, J. Anthony Wilson, Timothy S. Coalson, Bruce Fischl, Jesper L. Andersson, Junqian Xu, et al. 2013. “The Minimal Preprocessing Pipelines for the Human Connectome Project.” NeuroImage 80 (October):105–24.

Griffanti, Ludovica, Gholamreza Salimi-Khorshidi, Christian F. Beckmann, Edward J. Auerbach, Gwenaëlle Douaud, Claire E. Sexton, Enikő Zsoldos, et al. 2014. “ICA-Based Artefact Removal and Accelerated fMRI Acquisition for Improved Resting State Network Imaging.” NeuroImage 95 (July):232–47.

Hain, David, Tatiana Gallego-Flores, Michaela Klinkmann, Angeles Macias, Elena Ciirdaeva, Anja Arends, Christina Thum, et al. 2022. “Molecular Diversity and Evolution of Neuron Types in the Amniote Brain.” Science (New York, N.Y.) 377 (6610): eabp8202.

Heredia, Raúl, M. a. Angeles Real, Juan Suárez, Salvador Guirado, and José Carlos Dávila. 2002. “A Proposed Homology between the Reptilian Dorsomedial Thalamic Nucleus and the Mammalian Paraventricular Thalamic Nucleus.” Brain Research Bulletin 57 (3-4): 443–45.

Hua, Ruifang, Xu Wang, Xinfeng Chen, Xinxin Wang, Pengcheng Huang, Pengcheng Li, Wei Mei, and Haohong Li. 2018. “Calretinin Neurons in the Midline Thalamus Modulate Starvation-Induced Arousal.” Current Biology: CB 28 (24): 3948–59.e4.

Jakab, A., R. Blanc, E. L. Berényi, and G. Székely. 2012. “Generation of Individualized Thalamus Target Maps by Using Statistical Shape Models and Thalamocortical Tractography.” AJNR. American Journal of Neuroradiology 33 (11): 2110–16.

Jenkinson, Mark, Christian F. Beckmann, Timothy E. J. Behrens, Mark W. Woolrich, and Stephen M. Smith. 2012. “FSL.” NeuroImage 62 (2): 782–90.

Kark, Sarah M., Joren G. Adams, Mithra Sathishkumar, Steven J. Granger, Liv McMillan, Tallie Z. Baram, and Michael A. Yassa. 2022. “Why Do Mothers Never Stop Grieving for Their Deceased Children? Enduring Alterations of Brain Connectivity and Function.” Frontiers in Human Neuroscience 16 (September):925242.

Kark, Sarah M., Matthew T. Birnie, Tallie Z. Baram, and Michael A. Yassa. 2021. “Functional Connectivity of the Human Paraventricular Thalamic Nucleus: Insights From High Field Functional MRI.” Frontiers in Integrative Neuroscience 15 (April):662293.

Kochanski, Ryan B., Robert Dawe, Mehmet Kocak, and Sepehr Sani. 2018. “Identification of Stria Medullaris Fibers in the Massa Intermedia Using Diffusion Tensor Imaging.” World Neurosurgery 112 (April):e497–504.

Kooiker, Cassandra L., Matthew T. Birnie, and Tallie Z. Baram. 2021. “The Paraventricular Thalamus: A Potential Sensor and Integrator of Emotionally Salient Early-Life Experiences.” Frontiers in Behavioral Neuroscience 15 (May):673162.

Kooiker, Cassandra L., Yuncai Chen, Matthew T. Birnie, and Tallie Z. Baram. 2023. “Genetic Tagging Uncovers a Robust, Selective Activation of the Thalamic Paraventricular Nucleus by Adverse Experiences Early in Life.” Biological Psychiatry Global Open Science, January. 10.1016/j.bpsgos.2023.01.002.

Krauth, Axel, Remi Blanc, Alejandra Poveda, Daniel Jeanmonod, Anne Morel, and Gábor Székely. 2010. “A Mean Three-Dimensional Atlas of the Human Thalamus: Generation from Multiple Histological Data.” NeuroImage 49 (3): 2053–62.

Labouèbe, Gwenaël, Benjamin Boutrel, David Tarussio, and Bernard Thorens. 2016. “Glucose-Responsive Neurons of the Paraventricular Thalamus Control Sucrose-Seeking Behavior.” Nature Neuroscience 19 (8): 999–1002.

Leonard, Bianca T., Sarah M. Kark, Steven J. Granger, Joren G. Adams, Liv McMillan, and Michael A. Yassa. 2024. “Anhedonia Is Associated with Higher Functional Connectivity between the Nucleus Accumbens and Paraventricular Nucleus of Thalamus.” Journal of Affective Disorders 366 (December):1–7.

Li, Sa, and Gilbert J. Kirouac. 2012. “Sources of Inputs to the Anterior and Posterior Aspects of the Paraventricular Nucleus of the Thalamus.” Brain Structure & Function 217 (2): 257–73.

Mai, Jürgen K., Joseph Assheuer, and George Paxinos. 2004. Atlas of the Human Brain / Jürgen K. Mai, Joseph Assheuer, George Paxinos. 2nd ed. La Vergne, TN: Elsevier Academic Press.

Mai, Jürgen K., and Milan Majtanik. 2018. “Toward a Common Terminology for the Thalamus.” Frontiers in Neuroanatomy 12:114.

McGinty, Jacqueline F., and James M. Otis. 2020. “Heterogeneity in the Paraventricular Thalamus: The Traffic Light of Motivated Behaviors.” Frontiers in Behavioral Neuroscience 14 (October):590528.

McNally, Gavan P. 2021. “Motivational Competition and the Paraventricular Thalamus.” Neuroscience and Biobehavioral Reviews 125 (June):193–207.

Millan, E. Zayra, Zhiyi Ong, and Gavan P. McNally. 2017. “Paraventricular Thalamus: Gateway to Feeding, Appetitive Motivation, and Drug Addiction.” Progress in Brain Research 235 (September):113–37.

Morel, Anne. 2007. Stereotactic Atlas of the Human Thalamus and Basal Ganglia. 1st Edition. Boca Raton, FL: CRC Press. 10.3109/9781420016796.

Otis, James M., Manhua Zhu, Vijay M. K. Namboodiri, Cory A. Cook, Oksana Kosyk, Ana M. Matan, Rose Ying, et al. 2019. “Paraventricular Thalamus Projection Neurons Integrate Cortical and Hypothalamic Signals for Cue-Reward Processing.” Neuron 103 (3): 423–31.e4.

Percheron, G. 1973. “The Anatomy of the Arterial Supply of the Human Thalamus and Its Use for the Interpretation of the Thalamic Vascular Pathology.” Zeitschrift Für Neurologie 205 (1): 1–13.

Pfefferbaum, Adolf, Edith V. Sullivan, Natalie M. Zahr, Kilian M. Pohl, and Manojkumar Saranathan. 2023. “Multi-Atlas Thalamic Nuclei Segmentation on Standard T1-Weighed MRI with Application to Normal Aging.” Human Brain Mapping 44 (2): 612–28.

Reeders, Puck C., M. Vanessa Rivera Núñez, Robert P. Vertes, Aaron T. Mattfeld, and Timothy A. Allen. 2023. “Identifying the Midline Thalamus in Humans in Vivo.” Brain Structure & Function, January. 10.1007/s00429-022-02607-6.

Şahin, Mehmet Hakan, Abuzer Güngör, Oğuz Kağan Demirtaş, Çağrı Postuk, Zeynep Fırat, Gazanfer Ekinci, Hakan Hadi Kadıoğlu, and Uğur Türe. 2023. “Microsurgical and Fiber Tract Anatomy of the Interthalamic Adhesion.” Journal of Neurosurgery 139 (5): 1386–95.

Salimi-Khorshidi, Gholamreza, Gwenaëlle Douaud, Christian F. Beckmann, Matthew F. Glasser, Ludovica Griffanti, and Stephen M. Smith. 2014. “Automatic Denoising of Functional MRI Data: Combining Independent Component Analysis and Hierarchical Fusion of Classifiers.” NeuroImage 90 (April):449–68.

Saranathan, Manojkumar, Charles Iglehart, Martin Monti, Thomas Tourdias, and Brian Rutt. 2021. “In Vivo High-Resolution Structural MRI-Based Atlas of Human Thalamic Nuclei.” Scientific Data 8 (1): 275.

Schulmann, Anton, Ningping Feng, Pavan K. Auluck, Arghya Mukherjee, Ruchi Komal, Yan Leng, Claire Gao, et al. 2024. “A Conserved Cell-Type Gradient across the Human Mediodorsal and Paraventricular Thalamus.” bioRxivorg. 10.1101/2024.09.03.611112.

Shima, Yasuyuki, Henrik Skibbe, Yohei Sasagawa, Noriko Fujimori, Yoshimi Iwayama, Ayako Isomura-Matoba, Minoru Yano, et al. 2023. “Distinctiveness and Continuity in Transcriptome and Connectivity in the Anterior-Posterior Axis of the Paraventricular Nucleus of the Thalamus.” Cell Reports 42 (10): 113309.

Smith, Stephen M., and Thomas E. Nichols. 2009. “Threshold-Free Cluster Enhancement: Addressing Problems of Smoothing, Threshold Dependence and Localisation in Cluster Inference.” NeuroImage 44 (1): 83–98.

Toncray, J. E., and W. J. S. Krieg. 1946. “The Nuclei of the Human Thalamus; a Comparative Approach.” The Journal of Comparative Neurology 85 (3): 421–59.

Uroz, Victoria, Lucía Prensa, and José Manuel Giménez-Amaya. 2004. “Chemical Anatomy of the Human Paraventricular Thalamic Nucleus.” Synapse 51 (3): 173–85.

Van Buren, John M., and Rosemary C. Borke. 1973. Variations and Connections of the Human Thalamus: 1 the Nuclei and Cerebral Connections of the Human Thalamus. 2 Variations of the Human Diencephalon. PDF. Berlin, Germany: Springer. 10.1007/978-3-642-88594-5.

Vidal, Julie P., Kévin Rachita, Anaïs Servais, Patrice Péran, Jérémie Pariente, Fabrice Bonneville, Jean-François Albucher, Lola Danet, and Emmanuel J. Barbeau. 2024. “Exploring the Impact of the Interthalamic Adhesion on Human Cognition: Insights from Healthy Subjects and Thalamic Stroke Patients.” Journal of Neurology, July. 10.1007/s00415-024-12566-z.

Whitfield-Gabrieli, Susan, and Alfonso Nieto-Castanon. 2012. “Conn: A Functional Connectivity Toolbox for Correlated and Anticorrelated Brain Networks.” Brain Connectivity 2 (3): 125–41.

Winsky, L., P. Montpied, R. Arai, B. M. Martin, and D. M. Jacobowitz. 1992. “Calretinin Distribution in the Thalamus of the Rat: Immunohistochemical and in Situ Hybridization Histochemical Analyses.” Neuroscience 50 (1): 181–96.

Ye, Qiying, Jeremiah Nunez, and Xiaobing Zhang. 2022. “Oxytocin Receptor-Expressing Neurons in the Paraventricular Thalamus Regulate Feeding Motivation through Excitatory Projections to the Nucleus Accumbens Core.” The Journal of Neuroscience: The Official Journal of the Society for Neuroscience 42 (19): 3949–64.

