## Supplemental Figure 1 for "A combined neuroanatomy, ex vivo imaging and immunohistochemistry defined MRI mask for the human paraventricular nucleus of the thalamus"

### Supplementary Figures

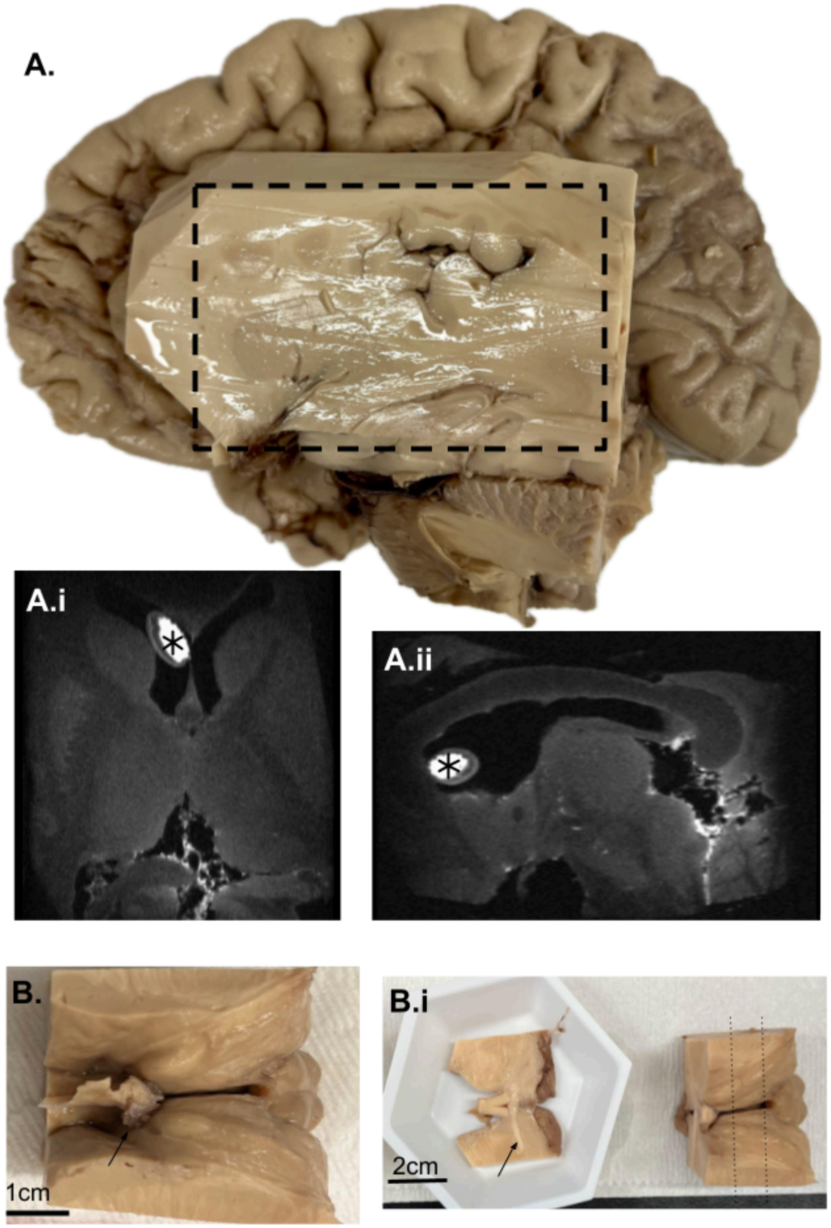


**Supplemental Figure 1: Tissue dissections.** (A) Cortex was dissected away from each hemisphere, leaving the ventricular area overlying the thalamus intact aside from a small opening to allow release of ventricular air bubbles, filling with fluorinert, and placement of a vitamin E tablet (asterisk) for right-left orientation. Transverse (i) and sagittal (ii) MRI planes display the extent of the tissue block dissection indicated on A with a dotted line. (B) Following MRI acquisition, the ventricle was dissected away to reveal the bilateral thalamus and facilitate tissue processing. Beginning at the anterior commissure (arrow), 1.5 cm slabs (B.i) were cut, placed in trays, embedded in OCT, and snap-frozen for cryosectioning.
